# Diverse tendencies in codon usage evolution of SARS-CoV-2 genes

**DOI:** 10.1101/2025.05.30.657095

**Authors:** Paweł Błażej, Dorota Mackiewicz, Paweł Mackiewicz

**Affiliations:** Department of Bioinformatics and Genomics, Faculty of Biotechnology, University of Wrocław, ul. Joliot-Curie 14a, 50-383 Wrocław, Poland

**Keywords:** coronavirus, codon usage, adaptation, optimization, translation, SARS-CoV-2

## Abstract

The dynamic evolution of SARS-CoV-2 virus since the COVID-19 outbreak in late 2019 has raised questions about potential evolutionary trends in protein-coding sequences and their adaptation to the human host. To address this, we compiled a dataset of 94,571 complete genomes with known collection dates, spanning from January 2020 to October 2024. Using a novel representation of codon usage, we recoded SARS-CoV-2 protein-coding sequences to strings of labels reflecting the human synonymous codon usage. Our analysis reveals different evolutionary pathways in the codon usage between structural, non-structural and accessory protein-coding sequences from the coronavirus. The genes coding for structural proteins tend to exhibit a less optimal adaptation to the human codon usage, whereas open reading frames *ORF1a* and *ORF1ab* encoding non-structural proteins show an opposite trend. The sequences for the accessory proteins demonstrated a variable tendency to change the codon preferences. The evolution of the more optimal codon usage in *ORF1a* and *ORF1ab* sequences can be associated with a higher speed and efficiency of translation of the coded polyproteins. Following their cleavage, the products play important roles in viral replication and transcription. Thus, the adaptation of their codons can increase the virus’ proliferation. In contrast, alterations in codon usage within structural protein-coding sequences may be associated with changes in their less accurate translation and folding during the synthesis, which can provide an advantage in evading the host immune response. The results show that codon usage adaptations to the human host differ based on the gene type and function, reflecting a balance between their conflicting evolutionary pressures. Our findings on variations in codon usage among coronavirus genes provide valuable insights that can aid in developing new strategies for the optimization of codons in vaccine mRNA and DNA for emerging strains.

## Introduction

Severe acute respiratory syndrome coronavirus 2 (SARS-CoV-2), the etiological agent responsible for the COVID-19 pandemic, emerged in Wuhan, China, in late 2019 (1). SARS-CoV-2 is classified as a betacoronavirus and shares a phylogenetic relationship with SARS-CoV and MERS-CoV but demonstrates distinct epidemiological and clinical characteristics (2).

The SARS-CoV-2 genome consists of a positive-sense, single-stranded RNA molecule of approximately 30 kb (3). The typical genome organization includes 14 open reading frames (ORFs), which encode 31 proteins. Among them, there are two large polyproteins (encoded by *ORF1a* and *ORF1ab*), four structural proteins and several accessory proteins (4, 5). The polyproteins are translated directly from the viral genomic RNA and subsequently cleaved into 16 nonstructural proteins, which perform essential functions in viral replication, transcription, RNA modification, protein processing as well as suppressing host gene expression and immune response.

To the structural proteins belong: surface glycoprotein known as Spike protein (S), envelope glycoprotein (E), membrane glycoprotein (M) and nucleocapsid glycoprotein (N) (6). The S protein is the key to host cell entry. It binds to the ACE2 receptor on the cell surface, triggering fusion of the viral and cellular membranes, allowing the virus to enter the cell. The E protein is a small protein involved in viral assembly, budding and release. It also acts as an ion channel and contributes to the pathogenicity of the virus, whereas the M protein is the most abundant structural protein. It helps maintain the virion’s shape and stability, interacts with other structural proteins and plays a crucial role in viral assembly. The N protein binds to the viral RNA genome, forming the ribonucleoprotein complex. It protects the viral RNA and is involved in viral replication and transcription. Products of other ORFs play important roles mainly in modulating host immune responses and facilitating viral replication.

After binding the Spike protein to the host cell receptor ACE2, the S protein undergoes conformational changes, allowing the viral envelope to fuse with the host cell membrane, releasing the viral genome into the cytoplasm (7). Since the viral RNA acts as mRNA, it is translated by host ribosomes into the two large polyproteins (8, 9). After their cleavage by viral proteases, they form a replication-transcription complex, which synthesizes new viral genomes through RNA-dependent RNA polymerase. Other proteins, structural and accessory ones, are produced by discontinuous transcription of subgenomic mRNAs and subsequent translation. The newly synthesized genomic RNA and N proteins interact with E and M proteins in the ER-Golgi intermediate compartment to form new virions, which are released from infected cells via exocytosis.

The viral strategy involves efficient replication and the production of vast array of proteins, which are then assembled into numerous virions. Thus, we can expect that the evolution of mechanisms accelerating these processes should be favoured. One of them can be changes in the synonymous codon usage, which can influence the speed and efficiency of translation (10–15). Many studies indicate that highly ex-pressed genes prefer codons, which are recognized by more numerous tRNA molecules. Since the SARS-CoV-2 utilizes the human host machinery, we can expect a relationship between the codon usage in the viral genes and the codon bias typical of *Homo sapiens* genes. The codon usage can also impact the speed and accuracy of translation elongation, causing changes in co-translational protein folding (15–21). It can also be important for modifying viral proteins that interact directly with the host’s immune system.

Certain analyses showed a general adaptation of viral to human codon usage (22, 23) or specifically to the upper respiratory tract and alveoli (24). However, other studies reported that strains with new mutations (25), all genes considered together (26, 27) or only genes for N and S proteins show a lower adaptation to the codon usage in the human host (28). Some authors noticed that the codon adaptation index (CAI) values calculated for the concatenated coding sequences of the coronavirus sequences have decreased over time with occasional fluctuations (29). However, the period considered in the study was rather short. The inconsistency in results reported by different authors may stem from the use of diverse methodologies and codon parameters, as well as the analysis of varied datasets. Therefore, we developed a new measure that recoded the codon usage in terms of the human host and studied a larger number of genomes collected from a wider evolution time of the coronavirus. Thanks to that, we could track temporal changes in protein-coding sequences over a longer time.

## Material and methods

### A. Dataset

We analyzed 94571 SARS-CoV-2 genomes downloaded from NCBI Virus database. We selected only complete genomes, isolated from human as well as possessing correctly annotated collection date and country. We studied protein-coding sequences from these genomes that were identified by VADR v1.4.2 - Viral Annotation DefineR (30) software. The reference model was vadr-models-sarscov2-1.3-2, which is based on NC_045512.2 RefSeq sequence obtained from isolate Wuhan-Hu-1 (31).

The downloaded set of genomes was a collection of genomes from 94 countries around the world. However, the proportion of genomes sequenced in the USA and the United Kingdom constituted over 88% of the dataset, i.e. 84324 genomes, hereby to avoid unbalanced sets, we decided to consider data only from these two countries. The collection dates for this filtered dataset spanned from January 2020 to October 2024.

### B. Codon usage recoding

We developed a novel measure termed codon block recoding (CBR), which maps the 64 codons to a set of ordered categorical labels reflecting the synonymous codon usage patterns. Specifically, CBR assigns each codon a label from 0 to 5, corresponding to its relative usage among synonymous codons encoding the same amino acid in protein-coding sequences of an organism. The most frequent codon in a synonymous block has assigned the label 0, whereas the least frequent codon has ascribed the label *n* − 1, where *n* is the number of codons in the block. Methionine and tryptophan, being represented by single codons, have the label 0. This mapping can be applied to any variant of genetic code and used for any genome. The labels {0, 1, 2, 3, 4, 5} are ordered categorical variables, which can be useful in qualitative studies. They are also easy to interpret and can be used, for example, to analyze the codons in groups characterized by distinct, high or low, relative usage. In this study, we applied human synonymous codon usage (HCU) to obtain human codon block recoding (HCBR), which is presented in Fig. 1. The table includes the canonical codon blocks encoding 20 amino acids and stop codons (*) with assigned relative frequencies of synonymous codons in human genes and assigned respective labels. The codon usage values were obtained from Codon Usage Database via python_codon_tables package (https://github.com/Edinburgh-Genome-Foundry/codonusage-tables/tree/master/python_codon_tables). The HCBR, or briefly the codon labels, were used to recode codons in individual coronavirus protein-coding sequences.

**Fig. 1.**
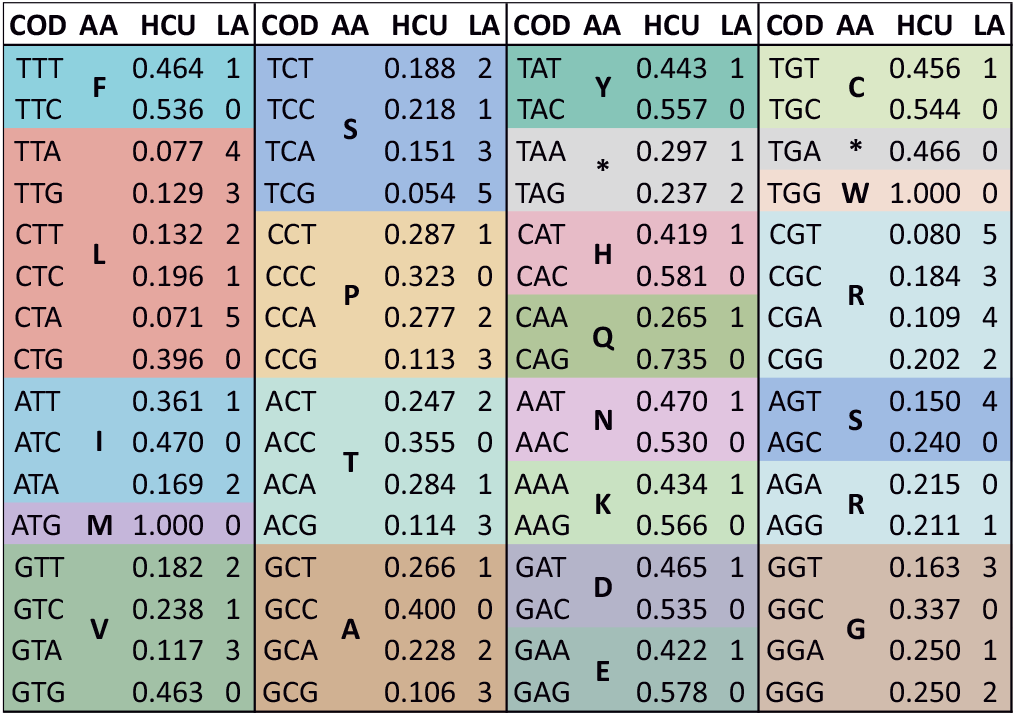
The human synonymous codon usage (HCU) table with assigned human codon block recoding (HCBR) labels. **Cod** - codon; **AA** - encoded amino acid; **HCU** - human synonymous codon usage; **LA** - assigned label corresponding to the usage

## Results

### C. Codon usage in a single sequence

The codon block recoding introduces a new representation of protein-coding sequences in terms of synonymous codon usage. Fig. 2 presents, as an example, the surface glycoprotein-coding sequence from the SARS-CoV-2 genome of isolate Wuhan-Hu-1, which was recoded according to HCBR. The individual codons have assigned labels corresponding the rank of synonymous codons according to their relative usage in the human protein-coding genes. Each dot represents a specific label of codon (y-axis) occupying its position in the sequence (x-axis). The codons labeled as 0, 1, 2 and 3 are predominant in the whole set. Some codons labeled as 4 and 5 are grouped in clusters in the sequences.

**Fig. 2.**
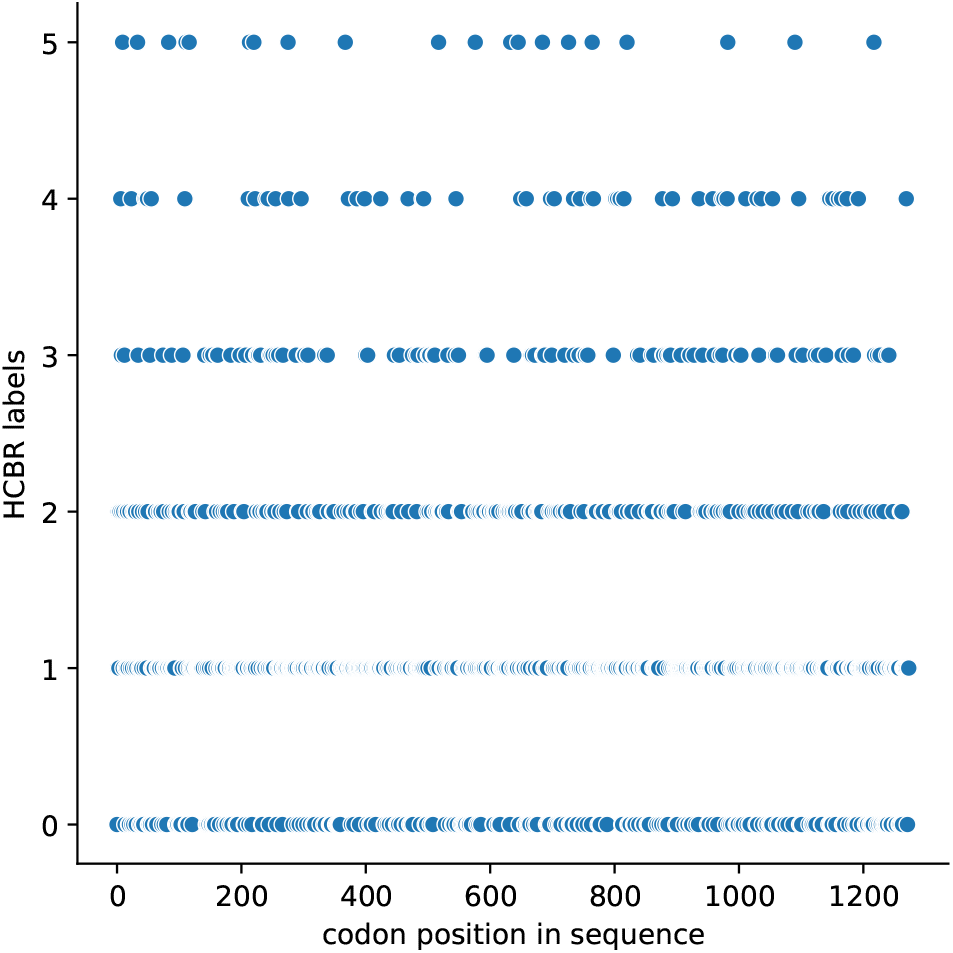
The surface glycoprotein-coding sequence from SARS-CoV-2 with codons recoded by its HCBR label.

The usage of recoded codons in the surface glycoprotein-coding sequence from the coronavirus differs from that in all human protein-coding genes (Fig. 3). The synonymous codons assigned to group 0, i.e. characterized by the highest relative usage in the human genes, are two times less frequent in the coronavirus sequence than in the genes, whereas those labeled 1 are more numerous. A slightly higher frequency is also for the less optimal codons with label 2, 3 and 4 in the surface protein gene than in the human genes. The frequency of codons 5 is comparable. The same tendencies for the first two codon types, we observed for other coronavirus protein-coding sequence with the exception to the envelope protein-coding sequence, in which the codons labeled 1 were also less frequent than in the human genes.

**Fig. 3.**
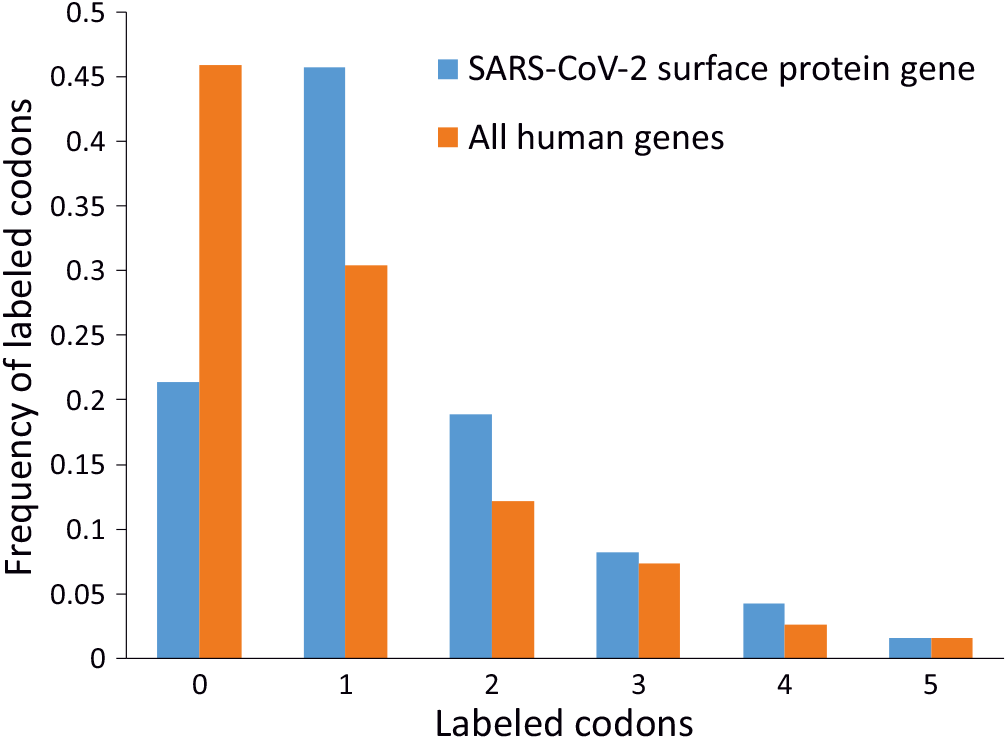
The distribution of codons labeled according to HCU for the surface glycoprotein-coding sequence from SARS-CoV-2 and all human genes.

Since we are interested in the codons commonly used in human protein-coding sequences, we calculated a combined frequency of 0 and 1 labeled codons, which measures the abundance of codons preferred in the human genes. The frequency of such codons in the example sequence is equal to 0.671, whereas in the human protein-coding sequences 0.763. Thus, the usage of these codons in the surface glycoprotein-coding sequence is lower than expected in human genes. Similarly, all other coronavirus gene sequences exhibit fewer of these codons than would be expected, especially the gene for envelope protein and *ORF10* (Table 1).

**Table 1.** The fraction of codons labeled 0 and 1 (0 + 1) calculated for SARS-CoV-2 protein-coding sequences from NC_045512.2 reference genome of isolate Wuhan-Hu-1.

| Gene | 0 + 1 | Gene | 0 + 1 |
| --- | --- | --- | --- |
| ORF1ab | 0.658 | ORF3a | 0.638 |
| ORF1a | 0.649 | ORF6 | 0.661 |
| surface | 0.671 | ORF7a | 0.705 |
| envelope | 0.513 | ORF7b | 0.705 |
| membrane | 0.659 | ORF8 | 0.680 |
| nucleocapsid | 0.695 | ORF10 | 0.590 |

### D. Changes of synonymous codon usage in time

The human codon block recoding was applied to track changes in the synonymous codon usage of individual protein-coding sequences from SARS-CoV-2 with time. We investigated eleven functional open reading frames (ORFs) or genes annotated in SARS-CoV-2 genomes: *ORF1ab, ORF1a*, gene for envelope protein, gene for membrane glycoprotein, gene for nucleocapsid phospoprotein, gene for surface glycoprotein as well as several accessory open reading frames: *ORF3a,ORF6,ORF7a,ORF8* and *ORF10*. The studied sequences were grouped according to coded structural, non-structural and accessory proteins, given their distinct roles in the viral life cycle. Following recoding, we calculated the frequency of codons most frequently used in human genes, i.e. labeled as 0 or 1, for each gene sequence. The values were aggregated monthly using arithmetic mean calculated for the sequences. This allowed us to describe the central point of available variants of protein-coding sequences in a month and find possible evolutionary trends in their codon usage in time.

#### Non-structural protein-coding sequences

Fig. 4 depicts changes in 0 + 1 codon frequencies over time for each non-structural protein-coding sequence, i.e. *ORF1a* and *ORF1ab* using the arithmetic mean calculated monthly. Since these two open reading frames overlap, on a long section, the changes in the frequency are very similar. The codons labeled as 0 + 1 were initially relatively rare from January 2020 to December 2021. From that to April 2022, we can observe a substantial shift towards higher frequencies of synonymous codons that are also the most commonly used in human genes. Next, their frequency remained at a similar relatively high level with a small decrease at the end of 2023 to the beginning of 2024. From that, the frequency gradually increased again. The global tendency, in the long run, indicates that these sequences optimized their synonymous codon usage to that in human protein genes.

**Fig. 4.**
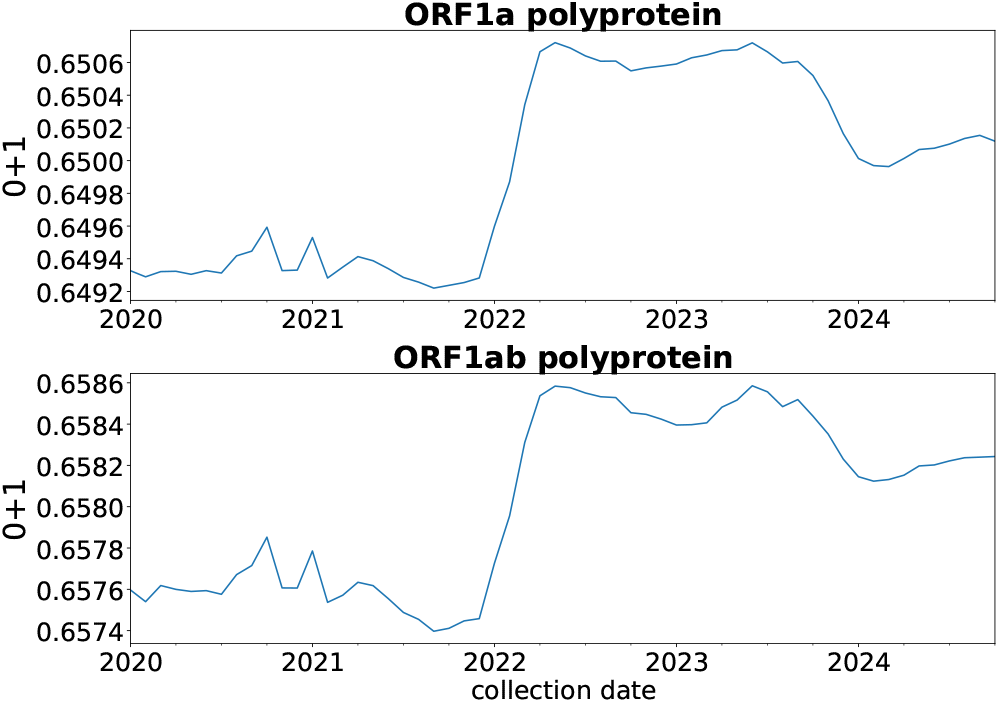
The changes in the frequency of codons labeled as 0 and 1 in SARS-CoV-2 non-structural protein-coding sequences from January 2020 to October 2024. The dynamic of changes was described by the arithmetic mean calculated monthly.

#### Structural protein-coding sequences

In contrast to the sequences coding for non-structural proteins, those encoding structural ones showed generally a decrease in the codons frequently used in the human host for a long time (Fig. 5). In all four sequences, the frequencies of 0 + 1 calculated at the end of 2024 are characterized by lower values in comparison those in January 2020. Thus, the structural protein-coding sequences tended to use less optimal codons in terms of HCU over time. Interestingly, the dynamic of changes differs between the sequences. In the case of the surface glycoprotein-coding sequence, we observed a general trend to decrease 0 + 1 frequency with several local extrema: the local minimum in September 2021 and February 2023 as well as the local maximum in February 2022 and October 2023 (Fig. 5). These fluctuations suggest that the decrease in the contribution of 0 + 1 codons was not linear but was disturbed by occasional variations.

**Fig. 5.**
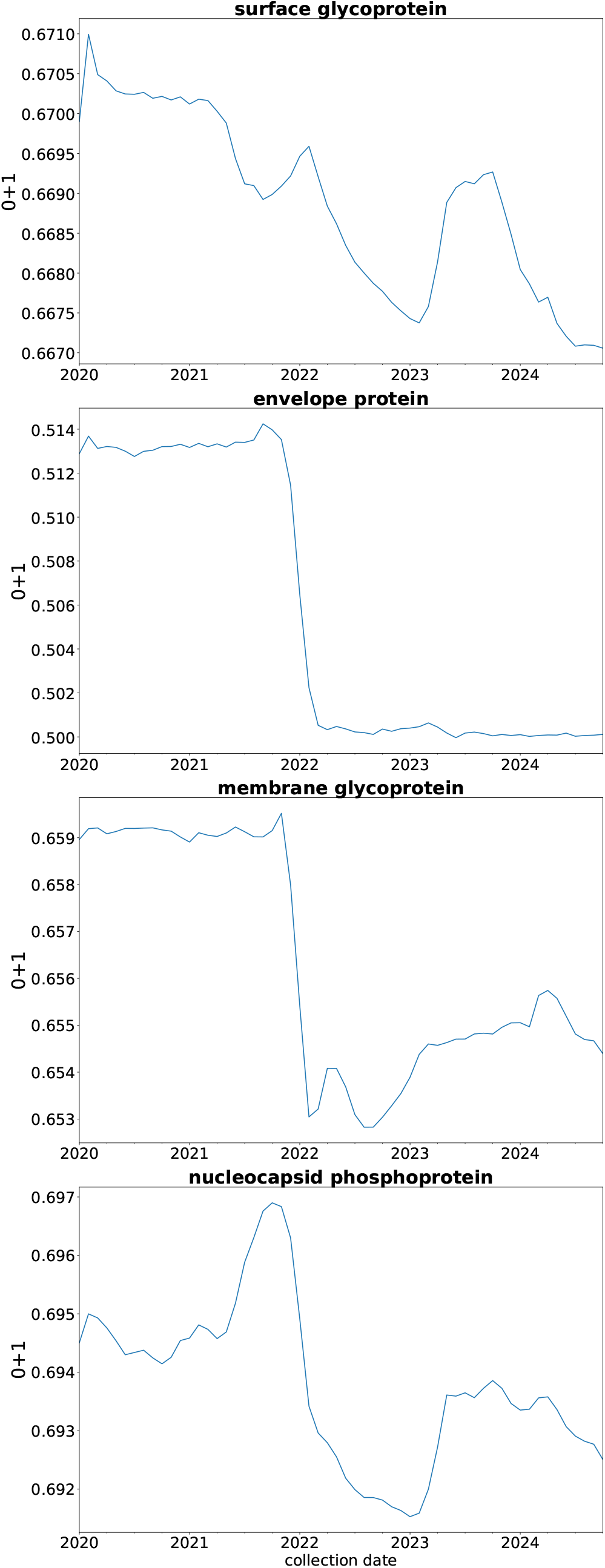
The changes in the frequency of codons labeled as 0 and 1 in SARS-CoV-2 structural protein-coding sequences from January 2020 to October 2024. The dynamic of changes was described by the arithmetic mean calculated monthly.

Three other structural protein-coding sequences demonstrated a characteristic decline from November 2021 up to: March 2022 (gene for envelope protein), February 2022 (gene for membrane glycoprotein) or August 2022 (gene for nucleocapsid phosphoprotein). All these curves have an inflection point at the border of years 2021 and 2022. The genes encoding the envelope protein and membrane glycoprotein exhibited a consistently high frequency of codons 0 and 1 up to November 2021. In contrast, the frequency of these codons in the nucleocapsid phosphoprotein gene fluctuated, peaking in October 2021. After the drastic decrease, the course of the curve for the codon frequency was stable through time but in the genes for membrane glycoprotein and nucleocapsid phosphoprotein it fluctuated reaching several local extrema.

#### Accessory protein-coding sequences

Other sequences coding for accessory proteins do not show consistent tendencies in the change of 0 + 1 codon frequencies (Fig. 6). However, we can notice a drop in this measure for *ORF6* sequence in the long term. Up to November 2021, the frequency was relatively high. Between November 2021 and April 2022 there was a drastic decrease with the minimum in March 2022 and a fast increase from April 2022 to September 2023. After a short stabilization at a relatively high level, the frequency diminished again from January 2023. Then, from June 2023 the frequency remained permanently low. *ORF3a, ORF7a* and *ORF8* showed a rather high and constant 0 + 1 frequency in time with an episodic quick decline and rise in various periods: November 2021-April 2022 (with the minimum in February 2022), February 2023-January 2024 (with the minimum in July 2023), February 2021-February 2022 (with the minimum in October 2021), respectively. In contrast to that, the frequency of codons 0 and 1 in *ORF10*, was rather low for the studied period but between May 2023 and May 2024, there was a sudden increase and decrease with the maximum in October 2023.

**Fig. 6.**
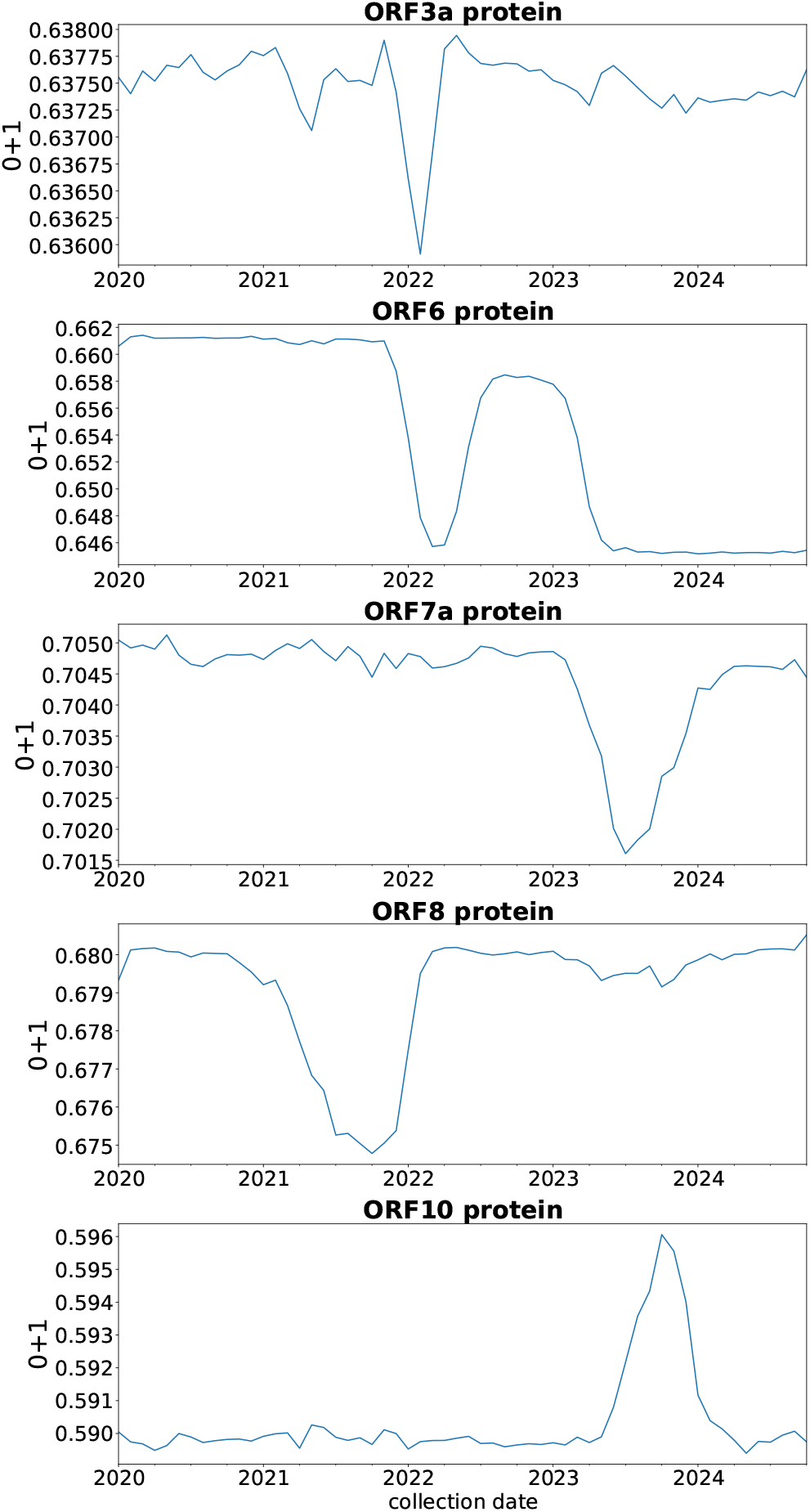
The changes in the frequency of codons labeled as 0 and 1 in SARS-CoV-2 non-structural protein-coding sequences from January 2020 to October 2024. The dynamic of changes was described by the arithmetic mean calculated monthly.

## Discussion

The recoding of codons according to the synonymous codon usage in human turned out useful in detecting changes in time in the codon preferences of protein-coding sequences from SARS-CoV-2 genomes. We studied the combined frequency of the two most widely used synonymous codons in human protein genes. Various groups of coronavirus gene sequences demonstrated different tendencies.

This frequency in *ORF1*_ab and *ORF_1a* increased as time progressed. Although the fraction of these codons never exceeded the expected value calculated for human codon usage, the increase in this frequency suggests a clear tendency for optimization of the viral codon usage with respect to human codon preferences. These ORFs encode polyproteins, which are cleavaged into smaller non-structural proteins. The enhanced adaptation of these sequences is likely linked to the critical role of their products in viral replication and the expression of other genes at the early stage of the virus infection cycle. Since the coronavirus utilizes the host translational machinery including tRNAs, the usage of codons also preferred in human genes can facilitate and speed up protein biosynthesis. This is in line with the general view that the synonymous codon bias can affect the rate and efficiency of translation. (10–15). Consequently, the more efficient translation leading to a higher yield of polyproteins and their products can accelerate viral proliferation and transmission. Our findings indicate that the codon usage of key viral proteins adapts over time to align with the host’s codon bias. In agreement with that, the general study of the codon usage in the coding sequences of 502 human-infecting viruses including SARS-CoV-2 found that the adaptation is visible in early viral proteins (24).

However, the structural protein-coding sequences exhibited a decreasing tendency in the frequency of the synonymous codons preferred by the human host. This suggests that these sequences tend to use less optimal codons in terms of HCU, which corresponds to the results by (28). The usage of poorly optimized codons may cause slower and more inaccurate translation elongation, which influences also the folding of synthetized proteins (15–21). Consequently, the overall production of viral proteins can be reduced. Moreover, the deviations from optimal translation rates as well as undesirable interactions between codons and non-cognate tRNAs can increase the number of various misfolded protein variants (32, 33). Since the structural proteins are exposed to the host immune system, their smaller number produced in non-optimal translation can be beneficial for the virus due to reduced recognition by the system. Moreover, the variable structure of epitopes can help to avoid the host response. Additionally, the rapid evolution of codon usage patterns over time can enhance the virus’s ability to adapt. In fact, we observed fluctuations in the usage of the optimal codons for genes encoding surface glycoprotein and nucleocapsid phosphoprotein eliciting a strong immune response (34, 35), which can be associated with changing the structure of these proteins with time.

The genes for accessory proteins demonstrated variable tendencies in the optimal codon usage. Due to their various function, it is difficult to find a general explanation for the changes in their codon frequencies. Nevertheless, the tendency to evolve and maintain the low frequency of the optimal codons observed in the structural and some accessory protein-coding sequences, can also associated with greater flexibility and adaptation to invade a broader range of hosts with different codon usage signatures (36, 37). In fact, it was found that SARS-CoV-2 can infect a range of mammalian species (38–40). It was also proposed that viruses with codon usage too similar to that of the host can be harmful to its cells due to the depletion of tRNA pool (41, 42). This can disrupt host translation and other cellular processes, potentially leading to over-exploitation of the host cell, which may ultimately limit the virus’s ability to multiply intensively.

Eight out of 11 studied protein-coding sequences revealed a drastic change in codon usage in a short time at the turn of the years 2021 and 2022. These changes correspond very well to the shift from Delta to Omicron variant (43–45). The Omicron was first identified on November 24, 2021 in South Africa and on December 1, 2021 in the United States. By the week ending December 25, it had become very quickly the dominant variant. The emergence of Omicron likely resulted from its high mutation rate, ability to evade immunity as well as the global connectivity and transmission that facilitated its rapid spread.

Using data filtered for USA and United Kingdom provided by (46), we also found that to the end of 2021 there was a significant increase in daily COVID-19 vaccine doses administered per million people, which culminated in December 2021. Vaccinations can also contribute to elevated selective pressure on the virus, favouring the emergence of mutations that enable it to evade immune responses. As a result, more viruses could replicate, thereby providing additional opportunities for mutations to occur (47, 48).

According to this dataset, daily new confirmed COVID-19 cases per million people started to grow from the middle of December 2021 and received the maximum in these countries in the middle of January 2022. The larger number of infected individuals provided also more opportunities for the virus to replicate, which in turn increases the probability of generating new mutations.

It is not inconceivable that the changes in codon usage observed in our studies resulted from the accumulation of new mutations and were associated with the phenomena mentioned above. Our findings also suggest that changes in codon usage, in terms of optimality for the human host, vary depending on the type and role of the genes. This reveals a trade-off between their competing evolutionary strategies.

Our results, highlighting variations in codon usage across different coronavirus protein-coding genes, provide valuable insights that can aid in designing effective attenuated vaccines for novel strains, thereby supporting efforts to combat the pandemic. This can be achieved by the influence of the codon (de)optimization in mRNA and DNA used in these vaccines.

## Notes

### Competing Interest Statement

The authors have declared no competing interest.

